# Increase in nutrient availability promotes success of invasive plants through increasing growth and decreasing anti-herbivory defenses

**DOI:** 10.1101/2021.10.18.464765

**Authors:** Liping Shan, Ayub M.O. Oduor, Wei Huang, Yanjie Liu

## Abstract

Invasive plant species often exhibit greater growth and lower anti-herbivory defense than native plant species. However, it remains unclear how nutrient enrichment of invaded habitats may interact with competition from resident native plants to affect growth and defense of invasive plants.

In a greenhouse experiment, we grew five congeneric pairs of invasive and native plant species under two levels of nutrient availability (low vs. high) that were fully crossed with simulated herbivory (clipping vs. no-clipping) and competition (alone vs. competition).

Invasive plants produced more gibberellic acid, and grew larger than native species. Nutrient enrichment caused a greater increase in total biomass of invasive plants than of native plants, especially in the absence of competition or without simulated herbivory treatment. Nutrient enrichment decreased leaf flavonoid contents of invasive plants under both simulated herbivory conditions, but increased flavonoid of native plants under simulated herbivory condition. Nutrient enrichment only decreased tannins production of invasive species under competition. For native species, it enhanced their tannins production under competition, but decreased the chemicals when growing alone.

The results indicate that the higher biomass production and lower flavonoids production in response to nutrient addition may lead to competitive advantage of invasive species than native species.

## 1. Introduction

Understanding the physiological and ecological processes underlying invasion success of alien plant species is an important topic in ecology (Jia *et al*., 2016; Reilly *et al*., 2020). Invasive plants commonly experience herbivory in their native ranges (Keane & Crawley, 2002; Wolfe, 2002), but because plant defense against herbivory incurs significant physiological and ecological costs (Cipollini *et al*., 2014), plants often have to trade off defense against growth and reproduction (Herms & Mattson, 1992). Therefore, theory predicts that alien plants that become successful invaders are those that have escaped from their own herbivores and re-allocated limited resources into greater growth and reproduction at the expense of defense (Keane & Crawley, 2002). In support it, several studies have reported that invasive plants interact with fewer herbivore species, and thus exhibit less defense and greater growth in the exotic range than in the native range (Colautti *et al*., 2004; Oduor *et al*., 2011; Meijer *et al*., 2016; Zhang *et al*., 2018). Therefore, alien plant species that become successful invaders may trade-off high growth and reproductive output with low investments in anti-herbivory defenses.

Observational studies have found that low-resource environments are generally less prone to invasion (Chytrý *et al*., 2008). Experimental studies also suggest that increased availability of resources for plant growth can confer invasive species with growth advantage over native species (D’Antonio & Vitousek, 1992; Bobbink *et al*., 1998; Davis *et al*., 2000; Tilman *et al*., 2001), becasue many native plant species are adapted to conditions of soil low-nutrient and water availability in their natural habitats (Bobbink *et al*., 1998; Dukes & Mooney, 1999). In fact, meta-analyses have found that nutrient enrichment is more beneficial to growth of invasive plant species than of native plant species (González *et al*., 2010; Liu *et al*., 2017). Following this logic, nutrient enrichment might also affect defense differently between invasive plant species and native plant species due to the trade-off between plant growth and defense (Herms & Mattson, 1992). However, how nutrient availability impacts growth-defense trade-offs of invasive and native plants remains little tested empirically.

Competition is important to determine plant invasion success (Levine *et al*., 2004; Petruzzella *et al*., 2020). On the one hand, strong competition from invasive plants often reduces diversity of native plant species, and results in mono-specific stands of invaders (Gaertner *et al*., 2009). Invasive plants exert strong competitive effects on native plants because invasive plants often have disproportionately higher demand for resources (Leishman & Thomson, 2005; Funk, 2013). Consequently, nutrient enrichment could confer invasive plants greater competitive advantage relative to native plants in communities (Seabloom *et al*., 2015). On the other hand, given that competition from other plants could create stressful environments, costs of plant defense against herbivory in such environment may also increase when competition is present (Herms & Mattson, 1992; Siemens *et al*., 2002). In other words, competition may amplify the growth-defense trade-offs of plants. However, it remains unclear whether competition affect trade-offs of invasive and native plants differently. Therefore, studies testing effects of tests of whether nutrient availability enrichment on growth-defense trade-offs of invasive and native plants, should also consider whether the plants grow alone or with competition.

Plant growth and defense are generally regulated by different types of hormones. For example, as the major hormones that stimulate plant growth and development (Ross & Reid, 2010), gibberellic acids (GA) stimulate seed germination, trigger stem elongation, leaf expansion, flowering and seed development (Yang *et al*., 2012; Gupta & Chakrabarty, 2013). However, expression of defense hormones can suppress expression of plant growth-promoting hormones, because these two type hormones often have negative cross-talks within the plants (Ross & Reid, 2010; Yang *et al*., 2012; Vos *et al*., 2015). For example, herbivory-induced production of a defense-regulating hormone jasmonic acid (JA) can constrain plant growth by antagonizing production of GA (Machado *et al*., 2017). Therefore, invasive plants that escape intense herbivores may produce high concentrations of growth-promoting hormones (e.g., GAs) and low concentrations of hormones that regulate anti-herbivore defenses (Liu *et al*., 2021). However, this prediction has not been tested empirically.

Here, we conducted a greenhouse experiment with five congeneric pairs of invasive and native plant species to test the following hypotheses: (i) Nutrient enrichment induces invasive plants to produce greater total biomass and lower concentrations of anti-herbivore defense compounds than native plants; (ii) Invasive plants express a lower concentration of a defense hormone JA and a higher concentration of a growth-promoting hormone GA.

## 2. Methods

### 2.1 Plant species

We used five congeneric pairs of native and invasive clonal plant species from three families that co-occur naturally in the field in China (**Table S1**). We raised plantlets/seedlings of the test plant species using seeds and asexual reproductive organs that were collected in the field (**Table S1**). For asexual species, we first selected intact rhizomes and stolons and cut them into single-node/bud fragments, and then cultivated the fragments in trays. For the sexually reproducing species, we directly sowed seeds in trays filled with potting soil (Pindstrup Plus, Pindstrup Mosebrug A/S, Denmark). The resultant plantlets/seedlings were then raised under uniform conditions for one month in a greenhouse (temperature: 22-28 □; natural lighting with an intensity of *c*. 75% of the light outdoors; and *c*. 60% relative humidity). We then selected similar-sized plantlets /seedlings of each species for use in the experiment described below.

### 2.2 Experimental set up

To test whether native and invasive plants differed in their responses to competition and herbivory at different levels of nutrient availability, we performed a fully-crossed factorial experiment with three factors: simulated herbivory (clipping vs. no-clipping), competition (alone vs. competition), and nutrient availability (low vs. high). Each treatment combination was replicated six times, which resulted in a total of 480 pots (i.e. 5 congeneric pairs of plants × 2 levels of invasion status per plant pair × 2 simulated herbivory levels × 2 competition levels × 2 nutrient treatment levels × 6 replicates). The plants were grown in 2.5-L circular plastic pots (top diameter × bottom diameter × height: 18.5 × 12.5 × 15 cm) that had been filled with a 1:1 mixture of sand and fine vermiculite. On 24^th^ March 2020, we transplanted 48 similar-sized individuals of each species into the center of the pot individually as target plants. For pots that received plant competition treatments, we planted three similar-sized seedlings of *Taraxacum mongolicum* around the target individual plants. We used *T. mongolicum* as a competitor because it commonly co-occurs with all the study species in various habitats in China (Chen *et al*., 2015). Plastic dishes were placed underneath each pot to hold water and nutrients that had been applied to the pots. All pots were randomly assigned to positions on four benches in a greenhouse, with a temperature range of 22 □ to 30 □ and natural light/darkness cycle.

We started the nutrient treatment on the second week following transplant. To impose low and high nutrient treatments, we added 0.2 and 1 g L^-1^ of a fertilizer solution (Peters Professional 20-20-20 General Purpose Fertilizer, Everris NA Inc., Dublin, OH, USA), respectively. Fifty milliliters of the respective nutrient solutions were supplied to the soil in the pots every week for 14 weeks. As all the experimental plant species were prone to herbivory by generalist above-ground herbivores in the natural habitats (Dang *et al*., 2012; Hu & Dong, 2019), we simulated above-ground herbivory effect on 17^th^ June 2020 (i.e., on the 12^th^ week following transplant). To remove *c*. 50% of leaf biomass of target plants as would happen under natural herbivory in the field, we clipped each leaf once by half across the midvein using a pair of scissors (for a similar approach, see Lurie *et al*., 2017; Kempel *et al*., 2020). The plants were watered once every day until harvest. The experiment was conducted in a greenhouse of Northeast Institute of Geography and Agroecology, Chinese Academy of Sciences (125°24’30”E, 43°59’49”N).

### 2.3 Harvest and measurements

On 27^h^ March 2020 (i.e. three days after transplanting), we measured initial height of each target individual plant. Two hours after clipping treatment (i.e. simulated herbivory application) on 17^th^ June 2020, we randomly selected three replicates of each target species under each treatment combination, and collected leaf samples for the measurement of GA3 (i.e. one of the principal gibberellic acids) and JA (Gupta & Chakrabarty, 2013; Camara *et al*., 2018). For the collection of leaves, we collected most recently matured leaves (the first fully expanded leaves, about 3-6 leaves below the shoot apex), froze them immediately in liquid nitrogen, and then stored the frozen samples at -80 □ until chemical analyses were performed as described below. Twelve days after the clipping treatment had been applied, we collected all leaf samples from the remaining three replicates of each target species under each treatment combination for the measurement of two groups of anti-herbivore defense compounds flavonoids and tannins. We used different replicates for plant hormones and defense compounds measurements, because we also cut leaves for plants under no-clipping condition when collecting samples for hormones measurements, which could induce changes of defense compounds. All leaves were dried at 40 □ to a constant biomass before being subjected to flavonoid and tannin analysis. We then harvested the left above-ground and below-ground biomass separately for each target plant immediately. The fresh biomass was then dried at 65 □ to constant weight and then weighed. As target or competitor plants died in six pots during the experiment, we harvested plants in 474 pots out of the original 480 pots. We calculated total biomass by summing the all above-ground and below-ground biomass. We also calculated the root mass fraction of each target plant as the ratio between the below-ground biomass and the total biomass.

For analysis of JA and GA3, all leaf samples were subjected to enzyme-linked immunosorbent assay (ELISA) procedure following previous studies (Dai *et al*., 2016). For analysis of flavonoid (quercetin, isoquercetin, quercetin glycoside, kaempferitrin, and kaempferol) and tannin (gallic acid, catechin, tannic acid, and ellagic acid) compounds, all leaf samples were firstly ground to a fine powder (< 0.25 mm) using a ball mill (MM400; Retsch, Haan, Germany), and then subjected to a high-performance liquid chromatography (HPLC) analysis (Wang *et al*., 2012). To quantify concentrations of individual flavonoids, 50 mg of ground leaf tissue samples were extracted with a 100% methanol-0.4% phosphoric acid (48:52, v:v) solution for 24 h. Next, we used 0.22-µm membrane to filter the extract, and then injected the filtrate (40 µL) into HPLC system for quantification. To quantify concentrations of tannins in plant leaves, 50 mg of ground leaf tissue samples were extracted in a 50% aqueous methanol solution for 30 min. Next, we used 0.45-µm membrane to filter the extract, and then injected the filtrate (40 µL) into HPLC system for quantification of individual tannins.

### 2.4 Statistical analyses

All statistical analyses were performed in R v4.0.2 (R Core Team, 2020). To test for main and interactive effects of species status, simulated herbivory, competition, and nutrient addition treatments on growth performance (i.e. total biomass production and root biomass allocation), concentrations of hormones (i.e. GA3 and JA) and chemical defense compounds (flavonoids and tannins) of target plants, we fitted linear mixed-effects models with the *lme* function in the *nlme* package (Pinheiro *et al*., 2020). In the models, total biomass, root mass fraction, leaf concentrations of GA3, JA, flavonoids, and tannins of target species were specified as response variables, while species status (native vs. invasive), simulated herbivory (clipping vs. no-clipping), competition (alone vs. competition), nutrient availability (low vs. high) treatments and their all interactions were specified as fixed-effect-independent variables. To meet the assumptions of normality of variance, total biomass and GA3 concentrations were square-root-transformed, while concentrations of flavonoids, tannins, and JA were natural-log-transformed. As initial size of plantlets/seedlings can influence the final growth performance of a species, we also added initial plant height as scaled natural-log-transformed covariates in the models for total biomass and root mass fraction.

To account for non-independence of replicates of the same species and for phylogenetic relatedness among target species, we also included target species nested within genus as random effects in all models. In addition, we also included reproduction modes as random factor in all models to account for the non-independence of replicates of the same reproduction modes. As the data did not fulfill homoscedasticity assumption, for the analyses of total biomass, root mass fraction and flavonoids, we included variance structures to allow for variance among species using the “*varIdent*” function in the R package “*nlme*” (Pinheiro *et al*., 2020). In the linear mixed-effect models described above, we assessed the significance of fix-effect independent variables (i.e., species status, competition, simulated herbivory, nutrient availability treatment and interactions among the four factors) using likelihood-ratio tests. The variance components were estimated using the restricted maximum-likelihood method of the full model (Zuur *et al*., 2009).

## 3. Results

### 3.1 Biomass production and allocation

Nutrient enrichment enhanced total biomass more for invasive species than for native species, in particular when plants grew alone (i.e., significant three-way interaction between species status, nutrient enrichment, and competition in Table 1; Fig. **1a**). Specially, the biomass increase of invasive species under competition was +127.0%, native species under competition +118.5%, invasive species growing alone +134.6%, and native species growing alone +124.3%. Moreover, nutrient enrichment caused a greater increase in total biomass of invasive plants than of native plants, especially in the absence of simulated herbivory condition (i.e. significant three-way interaction between species status, nutrient enrichment, and simulated herbivory in Table 1; Fig. **1b**). Specifically, the absolute biomass increase of invasive species under clipping treatment was +4.9 g, native species under clipping treatment +2.9 g, invasive species under no-clipping treatment +6.0 g, and native species under no-clipping treatment +3.3 g.

**Table 1.** Results of linear mixed-effect models that tested main and interactive effects of plant invasion status (invasive vs. native), nutrient enrichment (low vs. high), simulated herbivory (clipping vs. no-clipping), and competition (alone vs. competition) on growth performance of five congeneric pairs of plant species. Significant effects (*P* <0.05) are in bold, while marginally significant effects (0.05≤*P* <0.1) are underlined and in bold.

|  | Total biomass (g)<br>(square root transformed) |  | Root mass fraction<br>(logit- transformed) |  | Flavonoids (% dw)<br>(ln- transformed) |  | Tannins (% dw)<br>(ln- transformed) |  | GA3 (mg/g)<br>(square root- transformed) |  | JA (mg/g)<br>(ln- transformed) |  |
| --- | --- | --- | --- | --- | --- | --- | --- | --- | --- | --- | --- | --- |
| **Fixed effects** | $\chi^2(df = 1)$ | *p* | $\chi^2(df = 1)$ | *p* | $\chi^2(df = 1)$ | *p* | $\chi^2(df = 1)$ | *p* | $\chi^2(df = 1)$ | *p* | $\chi^2(df = 1)$ | *p* |
| Initial Height | 0.10 | 0.75 | 7.21 | **0.007** | - | - | - | - | - | - | - | - |
| Invasion status (S) | 5.66 | **0.017** | 3.77 | **0.052** | 1.15 | 0.284 | 1.57 | 0.211 | 4.69 | **0.030** | 3.44 | **0.064** |
| Nutrient enrichment (Nutr) | 304.52 | **<0.001** | 113.46 | **<0.001** | 5.22 | **0.022** | 0.07 | 0.786 | 0.36 | 0.547 | 2.74 | **0.098** |
| Clipping (Clip) | 15.56 | **<0.001** | 14.16 | **<0.001** | 2.99 | **0.084** | <0.01 | 0.980 | 0.73 | 0.392 | 0.16 | 0.692 |
| Competition (Comp) | 391.29 | **0.001** | 24.13 | **<0.001** | 7.04 | **0.008** | 4.47 | **0.035** | 1.32 | 0.251 | 1.50 | 0.221 |
| S × Nutr | 6.82 | **0.009** | 2.78 | **0.096** | 5.84 | **0.016** | 1.52 | 0.218 | 0.45 | 0.501 | 0.03 | 0.867 |
| S × Clip | 2.62 | 0.106 | 1.27 | 0.260 | <0.01 | 0.958 | 0.17 | 0.679 | 1.61 | 0.204 | 0.26 | 0.609 |
| S × Comp | 59.36 | **<0.001** | 0.19 | 0.660 | 1.61 | 0.205 | 1.31 | 0.252 | 0.07 | 0.797 | 0.11 | 0.746 |
| Nutr × Clip | 0.83 | 0.363 | -0.34 | 1.000 | 0.09 | 0.770 | 0.62 | 0.431 | 2.78 | **0.095** | 0.23 | 0.634 |
| Nutr × Comp | 49.56 | **<0.001** | <0.01 | 0.952 | 2.41 | 0.120 | 0.26 | 0.611 | 1.59 | 0.208 | 0.10 | 0.756 |
| Comp × Clip | 4.04 | **0.044** | 0.08 | 0.783 | <0.01 | 0.952 | 0.03 | 0.864 | 0.32 | 0.571 | 0.13 | 0.724 |
| S × Clip × Comp | 2.75 | **0.097** | 0.13 | 0.714 | 0.03 | 0.873 | 1.12 | 0.289 | 1.32 | 0.252 | 0.13 | 0.720 |
| S × Nutr × Clip | 4.74 | **0.030** | 0.05 | 0.818 | 5.72 | **0.017** | 0.52 | 0.473 | 1.34 | 0.247 | 0.75 | 0.386 |
| S × Nutr × Comp | 8.28 | **0.004** | 0.29 | 0.588 | 0.93 | 0.336 | 4.35 | **0.037** | 1.11 | 0.292 | 1.39 | 0.239 |
| Nutr × Clip × Comp | 0.29 | 0.591 | 1.65 | 0.199 | 2.27 | 0.132 | 1.46 | 0.227 | 0.49 | 0.483 | 0.37 | 0.542 |
| S × Nutr × Clip × Comp | 1.92 | 0.166 | 0.59 | 0.442 | 0.02 | 0.889 | 0.22 | 0.643 | 0.03 | 0.854 | 2.17 | 0.141 |
| **Random effects** | *SD* |  | *SD* |  | *SD* |  | *SD* |  | *SD* |  | *SD* |  |
| Genus | 0.555 |  | <0.001 |  | <0.001 |  | 0.531 |  | 0.107 |  | 0.196 |  |
| Species | 0.349* |  | 0.535* |  | 0.735* |  | <0.001 |  | 0.060 |  | 0.154 |  |
| Sowing | 0.004 |  | 0.531 |  | 0.226 |  | 0.264 |  | 0.168 |  | 0.242 |  |
| Residual | 0.288 |  | 0.641 |  | 1.296 |  | 0.586 |  | 0.209 |  | 0.473 |  |
| | $R^2_m$ | $R^2_c$ | $R^2_m$ | $R^2_c$ | $R^2_m$ | $R^2_c$ | $R^2_m$ | $R^2_c$ | $R^2_m$ | $R^2_c$ | $R^2_m$ | $R^2_c$ |
| $R^2$ of the model | 0.574 | 0.931 | 0.141 | 0.640 | 0.039 | 0.289 | 0.035 | 0.523 | 0.094 | 0.546 | 0.085 | 0.405 |

\* Standard deviations for individual species random effects for the saturated model are found in Table S2. $R^2_m$ : marginal $R^2$ , $R^2_c$ : conditional $R^2$ .

**Figure 1.**
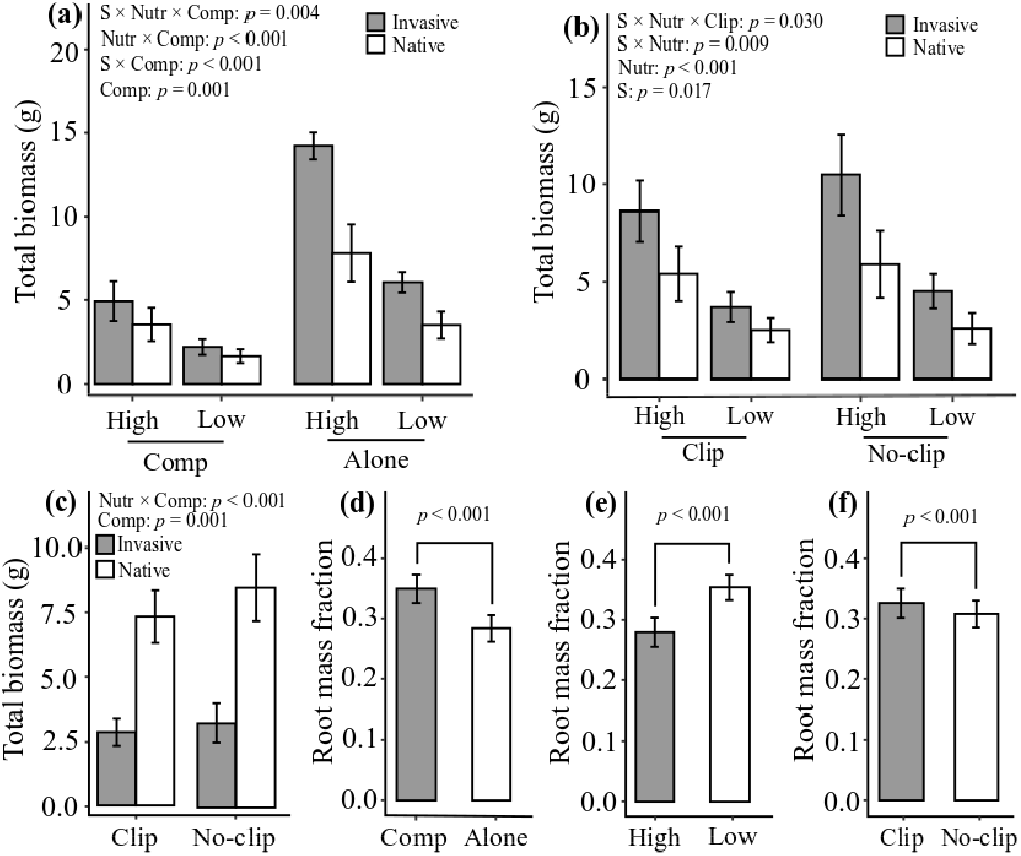
Mean (1 ± SE) biomass production and allocation of five congeneric pairs of invasive and native plant species that were grown in a common greenhouse condition. The panels show effects of: (**a**) three-way interaction between plant invasion status (invasive vs. native), nutrient enrichment (low vs. high), and competition (alone vs. competition [comp]) on total biomass; (**b**) three-way interaction between plant invasion status, nutrient enrichment, and simulated herbivory (clipping [clip] vs. no-clipping [no-clip]) on total biomass; (**c**) two-way interaction between competition and simulated herbivory on total biomass; (**d**) main effects of competition on plant root mass fraction; (**e**) main effects of nutrient enrichment on plant root mass fraction; (**f**) main effects of simulated herbivory on plant root mass fraction. The figure here only presents the significant main and interactive effects.

Total biomass production was also influenced significantly by two-way interactive effects of species status and nutrient enrichment, species status and competition, nutrient enrichment and competition, and competition and simulated herbivory (Table 1). When plants grew alone, simulated herbivory reduced the mean total biomass of target plants by -14.2%, and under competition, it reduced the total biomass by -13.0% (Fig. **1c**). Total biomass produced by the target plants was also influenced significantly by the main effects of plant invasion status, nutrient enrichment treatment, simulated herbivory, and competition (Table 1). Competition increased (Fig. **1d**), but nutrient enrichment decreased (Fig. **1e**) the proportion of total biomass that was allocated to the roots (i.e., root mass fraction). Simulated herbivory increased root mass fraction of the test plant species (Fig. **1f**).

### 3.2 Defense compounds

Simulated herbivory (i.e. leaf clipping) and nutrient enrichment had different effects on flavonoids concentrations in the leaves of invasive and native species (significant three-way interaction between plant invasion status, simulated herbivory, and nutrient availability treatment in Table 1; Fig. **2c**). Specially, for invasive plants, nutrient enrichment deceased concentrations of flavonoids similarly under both simulated herbivory treatments (clipping vs. no-clipping: -9.3% vs. -11.8%; Fig. **2c**). In contrast, for native plant species, nutrient enrichment increased flavonoid concentrations by +62.2 % under simulated herbivory (leaf clipping), whereas the reverse was true in the absence of simulated herbivory (no-clipping) treatment (−35.5%; Fig. **2c**). Flavonoids concentrations were also influenced significantly by a two-way interaction between plant invasion status and nutrient availability treatments, and by main effects of nutrient availability and competition treatments (Table 1). Competition increased the leaf concentrations of flavonoids by +27.8 % (Fig. **2a**) and of tannins by +13.2 % (Fig. **2b**). Leaf tannin concentrations were influenced by interactive effects of species status, competition, and nutrient enrichment (Table 1; Fig. **2d**). Specifically, for invasive plants, nutrient enrichment decreased tannin concentrations by -19.9% under competition, while it had little effect in the absence of competition (+0.7%; Fig. **2d**). For native species, nutrient enrichment enhanced tannin content under competition (+23.5%), whereas the reverse was true in absence of competition (Fig. **2d**).

**Figure 2.**
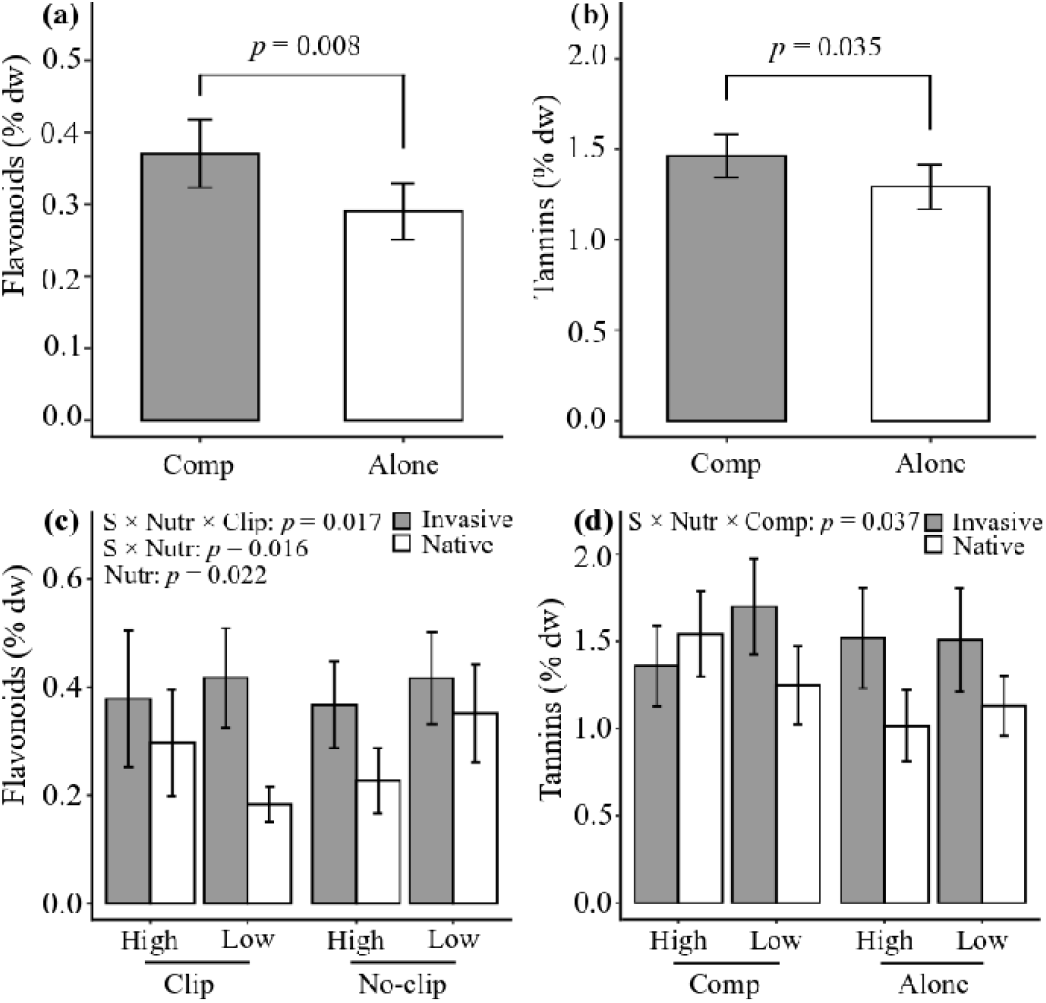
Mean (1 ± SE) concentrations of defense compounds (flavonoids and tannins) of five congeneric pairs of invasive and native plant species that were grown in a common greenhouse condition. The panels show effects of: (**a**) main effects of competition (alone vs. competition [comp]) on concentrations of flavonoids; **(b**) main effects of competition on concentrations of tannins; (**c**) three-way interaction between plant invasion status (invasive vs. native), nutrient enrichment (low vs. high), and simulated herbivory (clipping [clip] vs. no-clipping [no-clip]) on flavonoids concentrations; (**d**) three-way interaction between plant invasion status, competition and simulated herbivory on tannins concentrations. The figure here only presents the significant main and interactive effects.

### 3.3 Plant hormones

Invasive plants expressed higher (+24.9%) GA3 content in the leaves than native plants (Table 1; Fig. **3a**). The JA content in leaves was not significantly influenced by the separate effects of species status, nutrient enrichment, simulated herbivory, competition, and interactions among them (Table 1). Nevertheless, invasive plants tended to produce a greater amount of JA than native plants (*p* = 0.064; Fig. **3b**).

**Figure 3.**
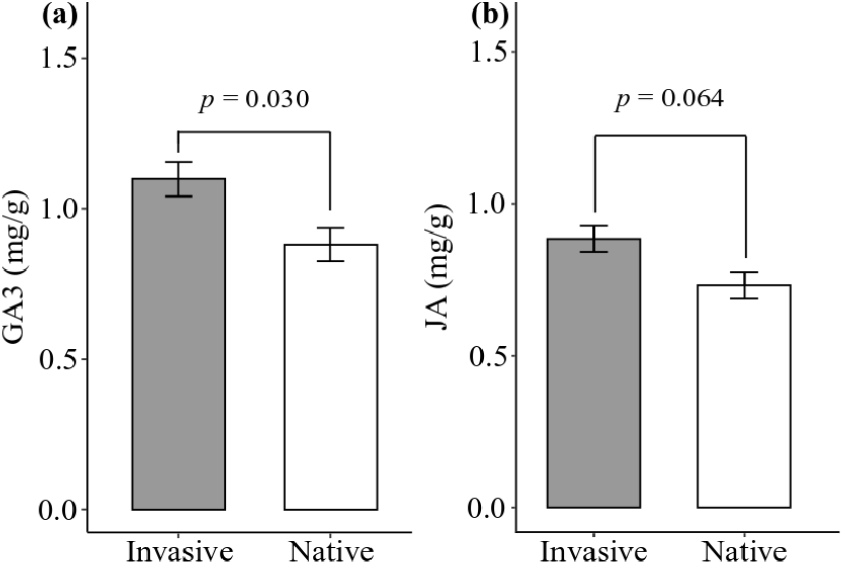
Mean (1 ± SE) concentrations of growth- and defense-enhancing hormones (GA3 and JA) of five congeneric pairs of invasive and native plant species that were grown in a common greenhouse condition. The panels show effects of main effect of plant invasion status (invasive vs. native) on (**a**) GA3 concentrations and (**b**) JA concentration.

## 4. Discussion

### 4.1 Biomass production and allocation

We found that nutrient enrichment enhanced total biomass production of invasive plant species more than it did for native plant species, more so in the absence of competition (Fig. **1a**). These results support those of other studies, which found that nutrient enrichment often promoted growth of invasive plant species over natives (Dawson *et al*., 2012; Parepa *et al*., 2013; Liu *et al*., 2017). Invasive plant species generally benefit more from an increase in resource availability than non-invasive species because invaders often have inherent fast growth strategies and the ability to rapidly exploit high-resource conditions (Dukes & Mooney, 1999; van Kleunen *et al*., 2010; Dawson *et al*., 2011). Moreover, high resource-use-efficiency can promote growth of some invasive species (Funk & Vitousek, 2007). Future studies may unravel which of these physiological mechanisms underlie the capacity of the current test invasive species to have greater total biomass than native species under nutrient enrichment. Overall, the present results support the idea that nutrient enrichment could enhance invasiveness of alien plant species that are already invasive.

Competition from a common native *T. mongolicum* lowered the beneficial effects of high nutrient enrichment for both native and invasive plants although the suppressive effect of the native competitor was higher on the invasive species (Fig. **1a**). These results suggest that the current invasive species are generally more sensitive to competition from *T. mongolicum* than the native plant species. As the native plants have co-existed with *T. mongolicum* longer than the invasive species have, it is likely that the native plant species are more strongly adapted to competition from *T. mongolicum* than the invasive species. Indeed, empirical studies show that native plants can evolve adaptation to competition from invasive plant species, and that native plants can also exert strong competitive effects on invasive plants (Oduor, 2013, 2021).

Our results complement those of other studies that tested the effects of nutrient enrichment on competitiveness of invasive plants, with variable outcomes. For example, Zhang *et al*. (2017) found that nutrient enrichment enhanced the competitive dominance of an invader *Alternanthera philoxeroides* over two natives *Oenanthe javanica* and *Iris pseudacorus*. In contrast, high nutrients diminished the competitiveness of an invasive herb *Hydrocotyle vulgaris* against a native plant community (Liu *et al*., 2016). Moreover, the invasive grass *Agrostis capillaris* suppressed growth of two co-occurring native grasses *Poa cita* and *Poa colensoi* in New Zealand regardless of nitrogen availability (Broadbent *et al*., 2018). The mix findings may because that these studies did not test the potential interactive effects of nutrient enrichment and herbivory on competitiveness of invasive and native plants. Our present study tested it, and found that nutrient enrichment caused a greater increase in total biomass of invasive plants than of native plants, especially in absence of simulated herbivory treatment (Fig. **1b)**. In other words, nutrient enrichment may act dependently of herbivory to affect competitiveness of invasive and native plants. Additional studies in other invasion systems may help to clarify the relative and combined effects of herbivory and nutrient enrichment on competitive interactions between invasive and native plant species.

Simulated herbivory caused a slightly greater decline in total biomass of the target plants in the absence of competition than in the presence of competition against *T. mongolicum* (Fig. **1c**), which suggests that simulated herbivory stimulated compensatory growth in the target plants. Herbivory can induce compensatory growth of the host plants specie through various mechanisms including increased rates of photosynthesis, increased growth rate, and increased allocation of biomass to the roots (Strauss & Agrawal, 1999; Stowe *et al*., 2000). We found that compensatory growth for invasive plants tended to be greater under competition than that in absence of competition (Fig. **S1**), which supports findings from other studies that simulated herbivory can stimulate compensatory growth in various invasive plants when grown in competition with other plants. For instance, leaf clipping of the invaders *Centaurea solstitialis* (Callaway *et al*., 2006) and *C. melitensis* (Callaway et al., 2008) and herbivory by a community of insects on invasive individuals of *Brassica nigra* (Oduor *et al*., 2013) enhanced growth of the host plants under interspecific competition. As the current focal plants had a greater root mass fraction under simulated herbivory and competition treatments (Fig. **1d**), it is likely that the focal plants deploy increased allocation of biomass to the roots as strategy to cope with simultaneous herbivory and competition. Future studies may unravel the other mechanisms of compensatory growth in the current study species.

### 4.2 Plant defense compounds

Invasive plants did not express a significantly lower concentration of the two classes of defense compounds tannins and flavonoids than native plants across the different treatment levels, which does not support a prediction of the idea that invaders should invest less in defense compounds (Keane & Crawley, 2002). As invaders escape from the specialist enemies but still face generalist enemies in the exotic range (Liu & Stiling, 2006; Meijer *et al*., 2016), it is plausible that invasive plants maintain a high concentrations of anti-herbivory chemicals to deter generalist herbivores. Indeed, Ni *et al* (2020) found that the invasive species *Alternanthera philoxeroides, Mikania micrantha* and *Wedelia trilobata* expressed significantly higher tannin concentrations than congeneric native species *A. sessilis, P. scandens* and *W. chinensis*. Similarly, total phenolic content of nine invasive plant species was 2.6-fold higher than that of nine native species in East Asia (Kim & Lee, 2011).

The finding that flavonoids and tannins concentrations were higher under competition treatment suggests that these metabolites may additionally have been involved in allelopathic interactions (Enge *et al*., 2012; Qi *et al*., 2020) between the target plants and the competitor species *T. mongolicum*. Invasive species produced greater concentrations of tannins than native species across the two nutrient treatments and in the absence of competition, although this trend was reversed by both high nutrient enrichment and competition (Fig. **2d**). This indicates that plant defense compounds tannins may play a greater allelopathic role in the native species than in the invasive species. In fact, flavonoids and tannins have been implicated in allelopathic interference in natural soils (Putnam & Duke, 1978; Weston & Mathesius, 2013), and it is likely to increase production of such allelopathic chemicals when stressed by other competitor plants (Lankau & Kliebenstein, 2009). This has been demonstrated for other groups of plant compounds. For example, *B. rapa* expressed higher concentrations of glucosinolate compounds when grown in interspecific competition (Siemens *et al*., 2002). In another study, the North American invader *B. nigra* increased production of glucosinolate compounds when grown in interspecific competition with three species (*Amsinckia menziesii, Malva parviflora*, and *Sonchus oleraceus*) (Lankau & Strauss, 2008). Future studies may test the relative allelopathic effects of tannins and flavonoids of the current test invasive and native plants against their natural competitors.

Nutrient treatment caused a decline in leaf flavonoid concentrations of invasive plants under both levels of simulated herbivory treatments, while it caused an increase in leaf flavonoid concentrations of native plants under simulated herbivory treatment and decreased the concentration under condition without simulated herbivory (Fig. **2c**). These results suggest that native plants rather than invasive plants may invest more in flavonoid-based defense against herbivores, and consequently grew slower than invasive plants in response to nutrient enrichment. On the other hand, under the condition without simulated herbivory, native plants would invest less in flavonoid-based defense against herbivores in response to nutrient addition. This is why native plants, in response to nutrient enrichment, increased growth stronger under condition without simulated herbivory than under simulated herbivory condition. Therefore, our study indicates that the increased investment in biomass and simultaneous decreased investment in flavonoid production by the focal invasive species in response to nutrient addition may facilitate their invasion success under scenarios of increased nutrient enrichment of invaded habitats.

### 4.3 Plant hormones

We found that invasive plants expressed a higher concentration of GA3 than native plants (Fig. **3a**). This supports our second hypothesis that invasive species often grow larger than native species because invaders produce greater amount of growth-promoting hormones (including GA3) than native species. However, contrary to our hypothesis, invasive plants did not express a lower concentration of JA than native plants (Fig. **3b**). Simulated herbivory induced a slightly higher amount of GA under high nutrient condition, but a lower amount of GA under low nutrient condition (Fig. **S2**). As the plants produced a lower amount of total biomass under simulated herbivory (Fig. **1b**), it is likely that an increase in GA stimulated a compensatory growth in the focal plants under high nutrient condition. Gibberellic acids can increase nutrient uptake, which is necessary for increased growth. For instance, exogenous application of GA on *Cicer arietinum* caused the plant to accumulate higher concentrations of the macronutrients nitrogen, phosphorus, and potassium (Rafique *et al*., 2021). Therefore, nutrient enrichment may have synergistic effects with GA on plant growth. The invasive plants expressed a slightly higher concentration of JA than native plants (Fig. **3b**). As the invasive plants produced higher concentrations of flavonoids under simulated herbivroy (especially under low nutrient condition), it is likely that the hormone stimulated a higher production of such defense compounds. In fact, exogenous spray of JA on plants has been shown to induce a high concentration of flavonoids (War *et al*., 2014). The higher concentrations of JA and the flavonoids content in invasive plants suggest that the plants may be more strongly defended against herbivores than native plants.

## Conclusion

Our results indicate that the higher expression of growth-related hormones could contribute the larger growth of invasive species than native species. Moreover, the lower investment of invasive species in the anti-herbivory chemicals when they grew under high nutrient conditions could lead to their stronger growth response to nutrient enrichment. Our study provides the evidence that higher investment of growth and lower investment of anti-herbivory chemicals, at least flavonoids in response to nutrient enrichment would lead to competitive advantage of invasive alien species than native species.

## Acknowledgement

We thank Xue Zhang, Lichao Wang, and Zaixing Ma for their practical assistance. YL acknowledges funding from the Chinese Academy of Sciences (Y9B7041001). AO acknowledges funding from the CAS President’s International Fellowship Initiative (2021VBB0004).

## Author contributions

YL conceived the idea and designed the experiment. LS performed the experiment. LS and YL analyzed the data. LS and AO wrote the first draft of the manuscript, with major inputs from YL and further inputs from WH.

## Data accessibility

Should the manuscript be accepted, the data supporting the results will be archived in Dryad and the data DOI will be included at the end of the article.

## Supporting information

**Table S1.**
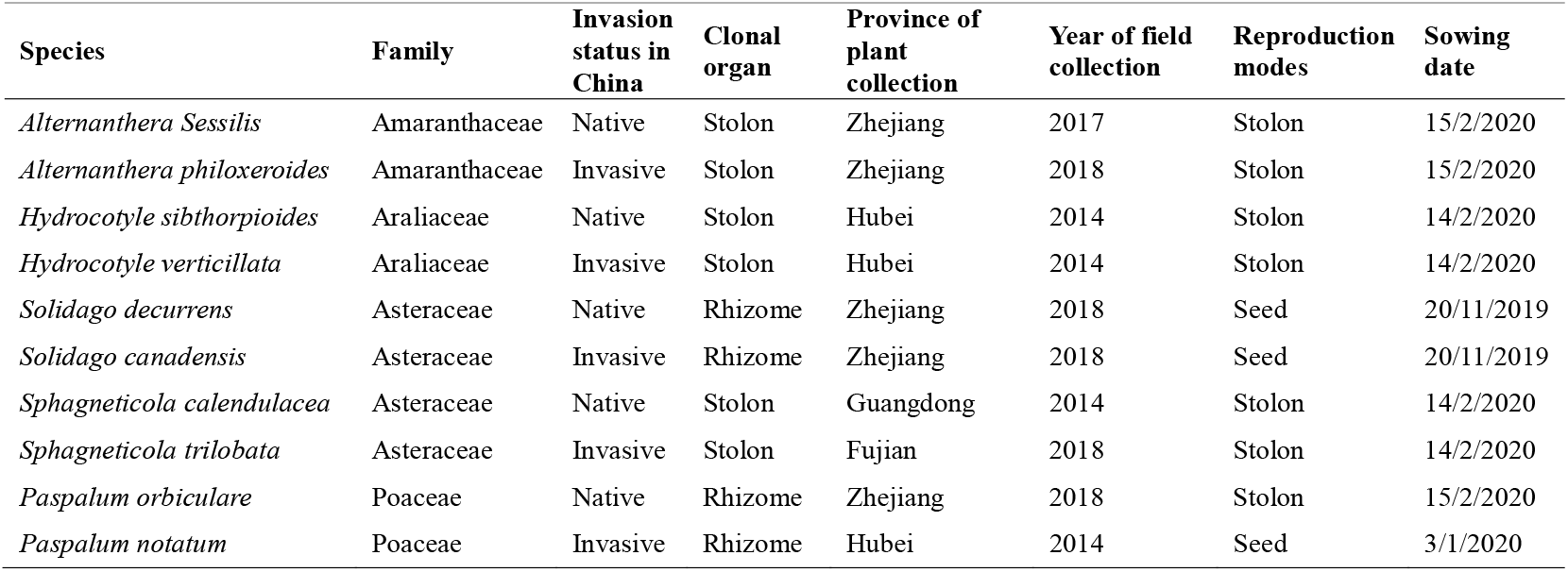
Information on the five congeneric pairs of invasive and native plant species that were used in the current experiment

| Species | Family | Invasion status in China | Clonal organ | Province of plant collection | Year of field collection | Reproduction modes | Sowing date |
| --- | --- | --- | --- | --- | --- | --- | --- |
| *Alternanthera Sessilis* | Amaranthaceae | Native | Stolon | Zhejiang | 2017 | Stolon | 15/2/2020 |
| *Alternanthera philoxeroides* | Amaranthaceae | Invasive | Stolon | Zhejiang | 2018 | Stolon | 15/2/2020 |
| *Hydrocotyle sibthorpioides* | Araliaceae | Native | Stolon | Hubei | 2014 | Stolon | 14/2/2020 |
| *Hydrocotyle verticillata* | Araliaceae | Invasive | Stolon | Hubei | 2014 | Stolon | 14/2/2020 |
| *Solidago decurrens* | Asteraceae | Native | Rhizome | Zhejiang | 2018 | Seed | 20/11/2019 |
| *Solidago canadensis* | Asteraceae | Invasive | Rhizome | Zhejiang | 2018 | Seed | 20/11/2019 |
| *Sphagneticola calendulacea* | Asteraceae | Native | Stolon | Guangdong | 2014 | Stolon | 14/2/2020 |
| *Sphagneticola trilobata* | Asteraceae | Invasive | Stolon | Fujian | 2018 | Stolon | 14/2/2020 |
| *Paspalum orbiculare* | Poaceae | Native | Rhizome | Zhejiang | 2018 | Stolon | 15/2/2020 |
| *Paspalum notatum* | Poaceae | Invasive | Rhizome | Hubei | 2014 | Seed | 3/1/2020 |

**Table S2.**
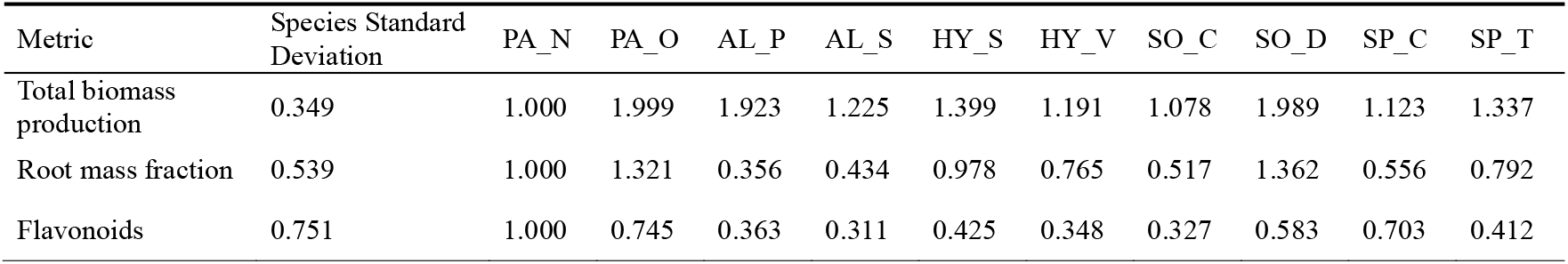
Standard deviations for individual species random effects for metrics analyzed with models with a Gaussian error distribution. The standard deviations given refer to the first species. For each species, these should be multiplied by the multiplication factors. The names of the species in the table are abbreviated using the first and second letters of the genus and the first letter of species epithet.

| Metric | Species Standard Deviation | PA_N | PA_O | AL_P | AL_S | HY_S | HY_V | SO_C | SO_D | SP_C | SP_T |
| --- | --- | --- | --- | --- | --- | --- | --- | --- | --- | --- | --- |
| Total biomass production | 0.349 | 1.000 | 1.999 | 1.923 | 1.225 | 1.399 | 1.191 | 1.078 | 1.989 | 1.123 | 1.337 |
| Root mass fraction | 0.539 | 1.000 | 1.321 | 0.356 | 0.434 | 0.978 | 0.765 | 0.517 | 1.362 | 0.556 | 0.792 |
| Flavonoids | 0.751 | 1.000 | 0.745 | 0.363 | 0.311 | 0.425 | 0.348 | 0.327 | 0.583 | 0.703 | 0.412 |

**Figure S1.**
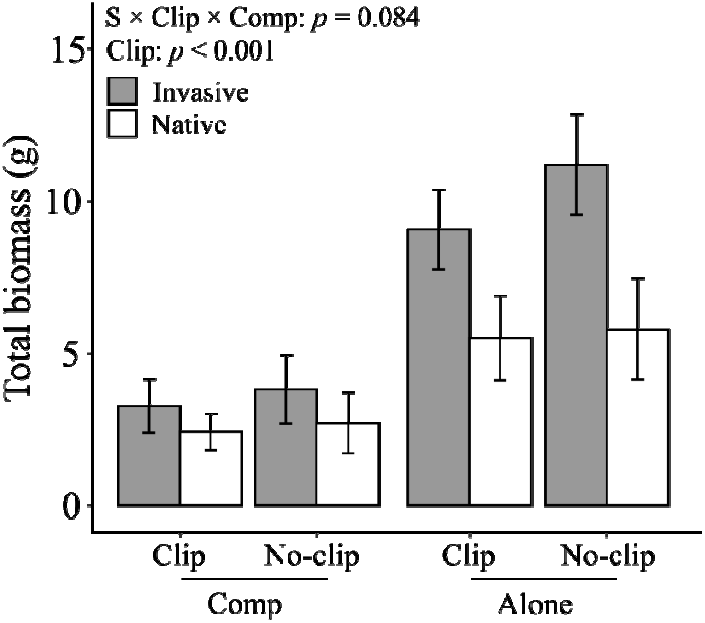
Mean (1± SE) total biomass of five congeneric pairs of invasive and native plant species that were grown in a common greenhouse condition. The panels show effects of three-way interaction between plant invasion status (invasive vs. native), competition (alone vs. competition [comp]), and simulated herbivory (clipping [clip] vs. no-clipping [no-clip]) on total biomass.

**Figure S2.**
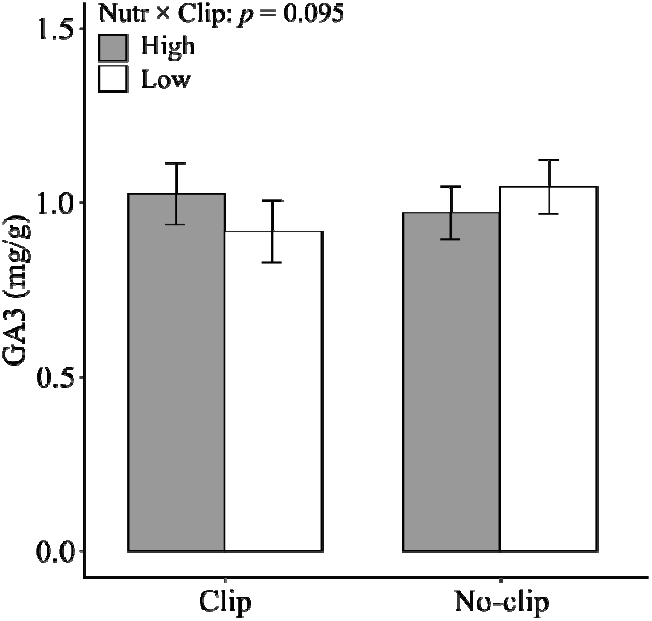
Mean (1± SE) concentrations of GA3 of five congeneric pairs of invasive and native plant species that were grown in a common greenhouse condition. The panels show effects of two-way interaction between nutrient enrichment (low vs. high) and simulated herbivory (clipping [clip] vs. no-clipping [no-clip]) on GA3 concentrations.

